## Supplementary Information for "Systematic conformation-to-phenotype mapping via limited deep-sequencing of proteins"

#### Supplementary Figures

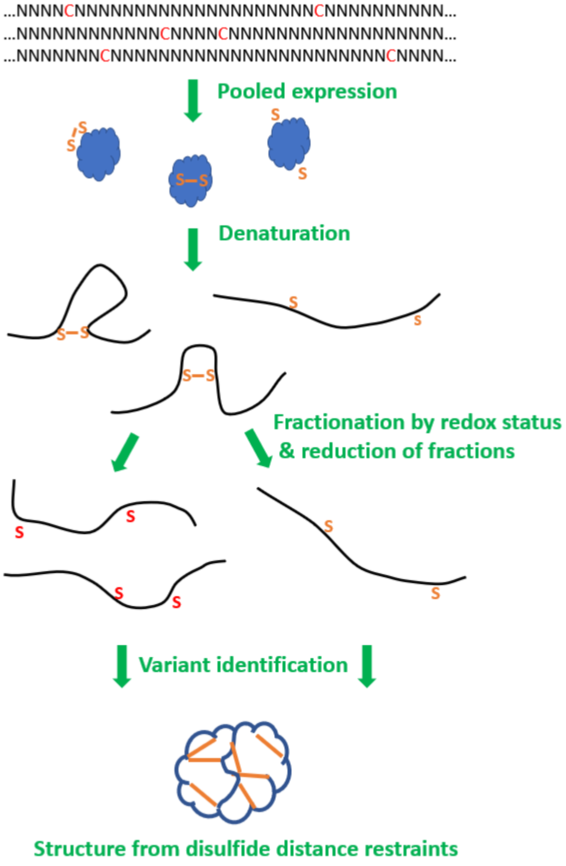


**Figure SI 1: A hypothetical illustration of protein structure determination by high-throughput disulfide scanning (HTDS).** The essence of the technique is to reduce the 3D problem of protein structure mapping to the 1D problem of protein sequencing. First, a pooled double-Cys scanning library is expressed, and as the proteins fold, pairs of Cys are brought sufficiently close together in the tertiary structure to form disulfides, while other pairs are separated and remain reduced. This effectively encodes the 3D structure of the protein in a 2D matrix of disulfide formation propensity as a function of the two sequence positions. The tertiary structure can then be denatured without loss of structural information, provided the disulfides remain intact. The library is then fractionated by redox status. This can be accomplished either by thiol affinity chromatography or – as in our study – by simply alkylating all free thiols at this step. This effectively encodes the 2D matrix of disulfide propensities in the 1D sequences of the protein variants. Disulfides can then be reduced without loss of structural information, provided the polypeptide backbones remain intact. The disulfide-forming variants are then identified by protein sequencing. In the current version of our method, this means cyanylation of the Cys thiols followed by aminolysis of the polypeptide backbone immediately upstream of the cyano-Cys sites. MS/MS then identifies and quantifies each double-Cys variant by measuring the middle fragment from this Cys-targeted backbone cleavage. Finally, a computational method is required to back-calculate the original 3D structure from the 2D matrix of disulfide bonding propensities.

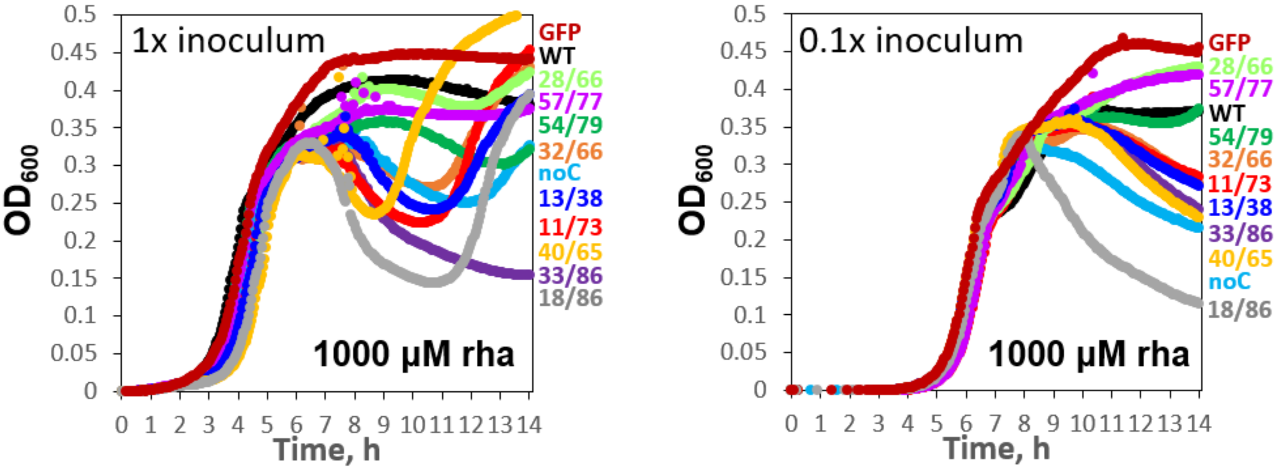


**Figure SI 2: Effect of inoculum size on the growth curve dynamics of HdeA variant-expressing Δ*hdea* BW25113 *E. coli*.** *Left*: the same plot as **Figure 2a**, shown for reference. *Right*: the equivalent experiment with 1/10 the inoculum size.

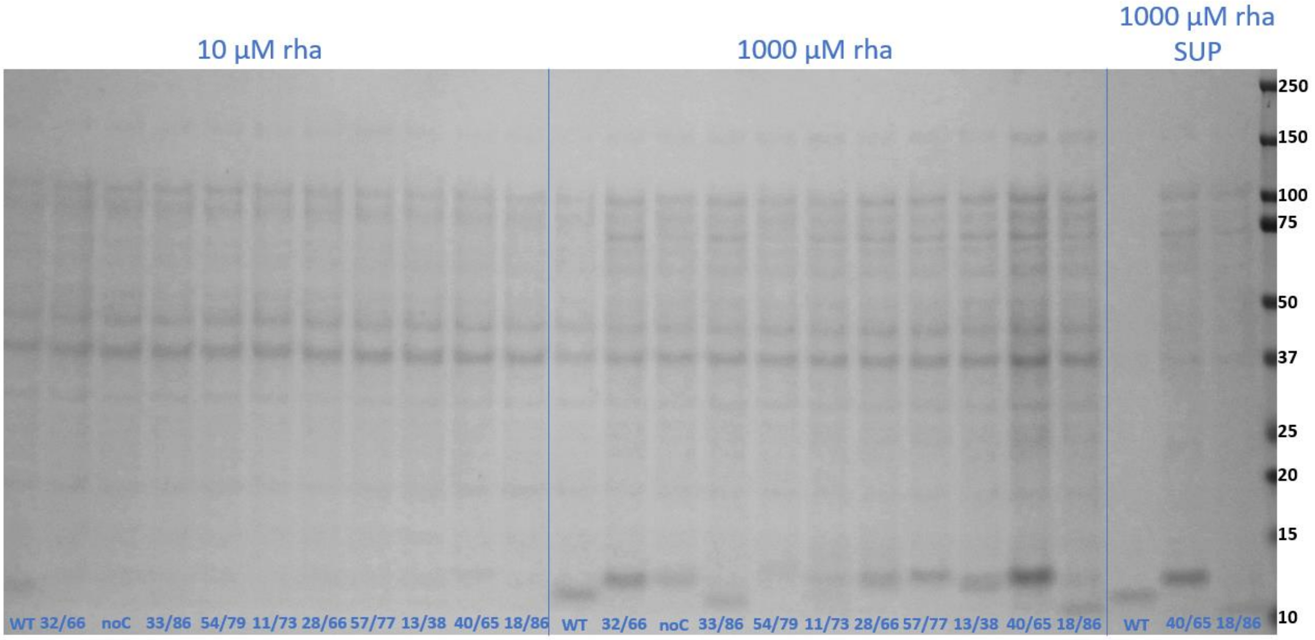


**Figure SI 3: Non-reducing SDS-PAGE of total end-point cultures grown and induced as in Figure 2b, but without using MOPS buffer in the LB medium during initial (pre-induction) overnight culture.** Expression levels were more variable under these conditions; at 10 µM rhamnose, only the WT was visibly expressed, but at 1000 µM rhamnose, all variants had visible HdeA bands consistent with their expected migration patterns.

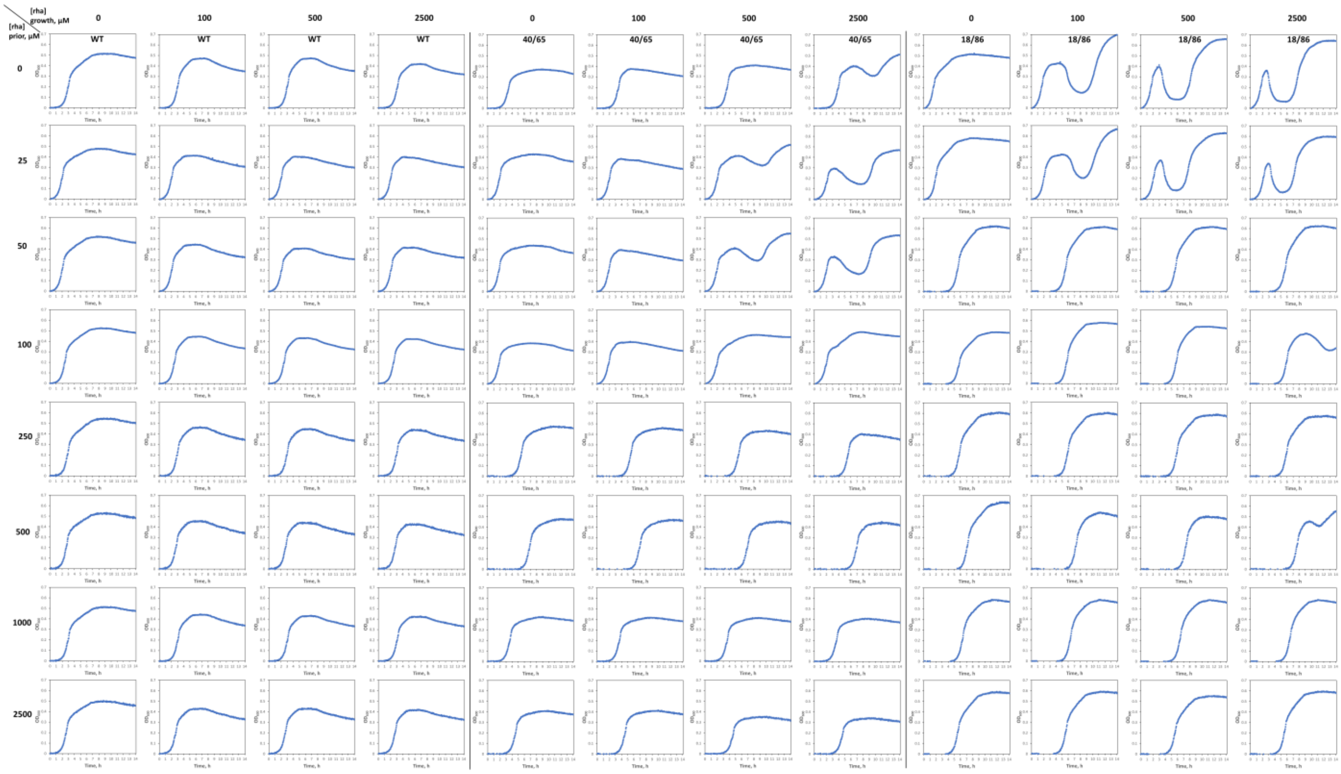


**Figure SI 4: *E. coli* cultures show persistent adaptation after recovery from cytotoxic HdeA variant expression.** Three variants were compared: WT HdeA (left four columns), variant 40/65 (middle four columns), and variant 18/86 (right four columns). Numbers from top to bottom indicate [inducer] during initial culture, from 0 to 2500 µM. Numbers across the top indicate [inducer] during a subsequent culture inoculated from the initial culture: 0, 100, 500, or 2500 µM for each variant. As expected, when the initial culture was done without inducer (*top row*), the growth curves of the two cytotoxic variants (but not of the WT) showed a strong die-off dependent on induction level, followed by recovery. However, when initial cultures contained inducer (*subsequent rows*), and the second culture was inoculated from the recovered initial cultures, then the die-off phenotype disappeared. However, these recovered cultures had visibly longer lag phases than the naïve cultures, suggesting an epigenetic memory that alters growth dynamics.

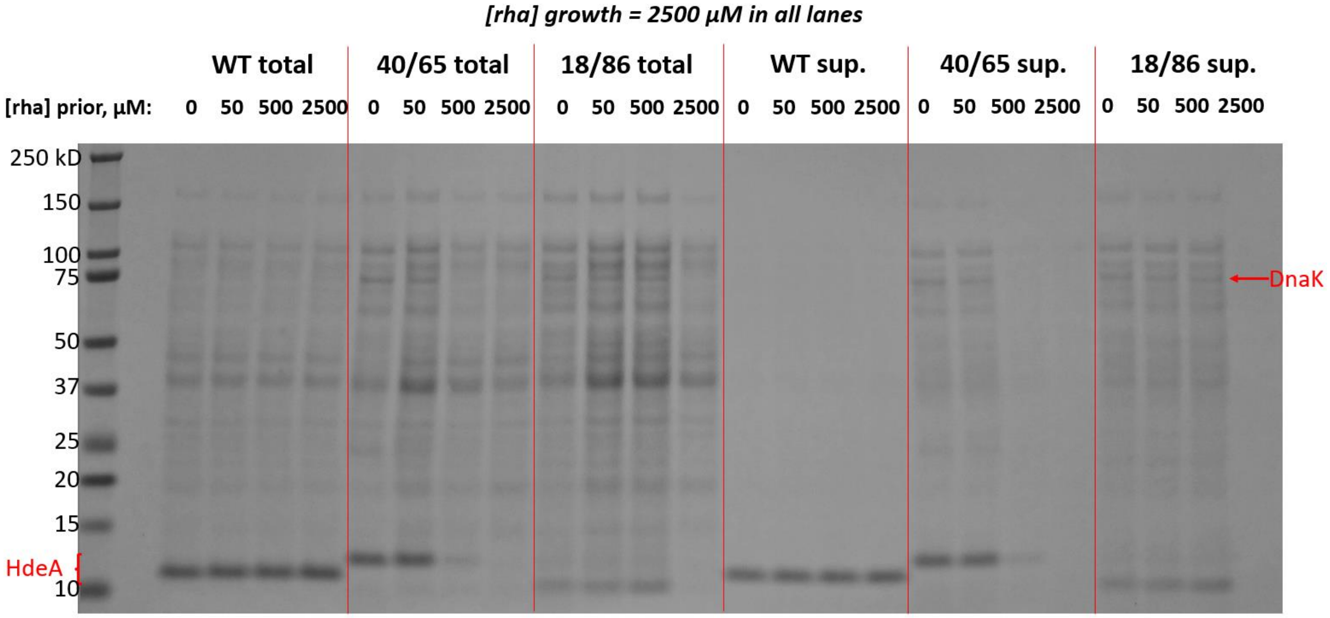


**Figure SI 5: Recovery from cytotoxicity caused by HdeA misfolding entails loss of HdeA expression.** Non-reducing SDS-PAGE of total cell culture and cell culture supernatants from the endpoints of the respective growth curves shown in columns 4, 8, and 12 of **Figure SI 3** (i.e., secondary cultures grown in the presence of 2500 µM rhamnose). Inducer concentrations during primary culture were as indicated: 0, 50, 500, or 2500 µM. WT HdeA cultures continued to express the protein ("WT total" lanes), but 40/65 and 18/86 cultures lost protein expression if the primary cultures contained high [rhamnose] ("40/65 total" and "18/86 total" lanes). Moreover, the overexpression of endogenous DnaK (*red arrow*) likewise diminished, as did cytolysis (the "sup." lanes).

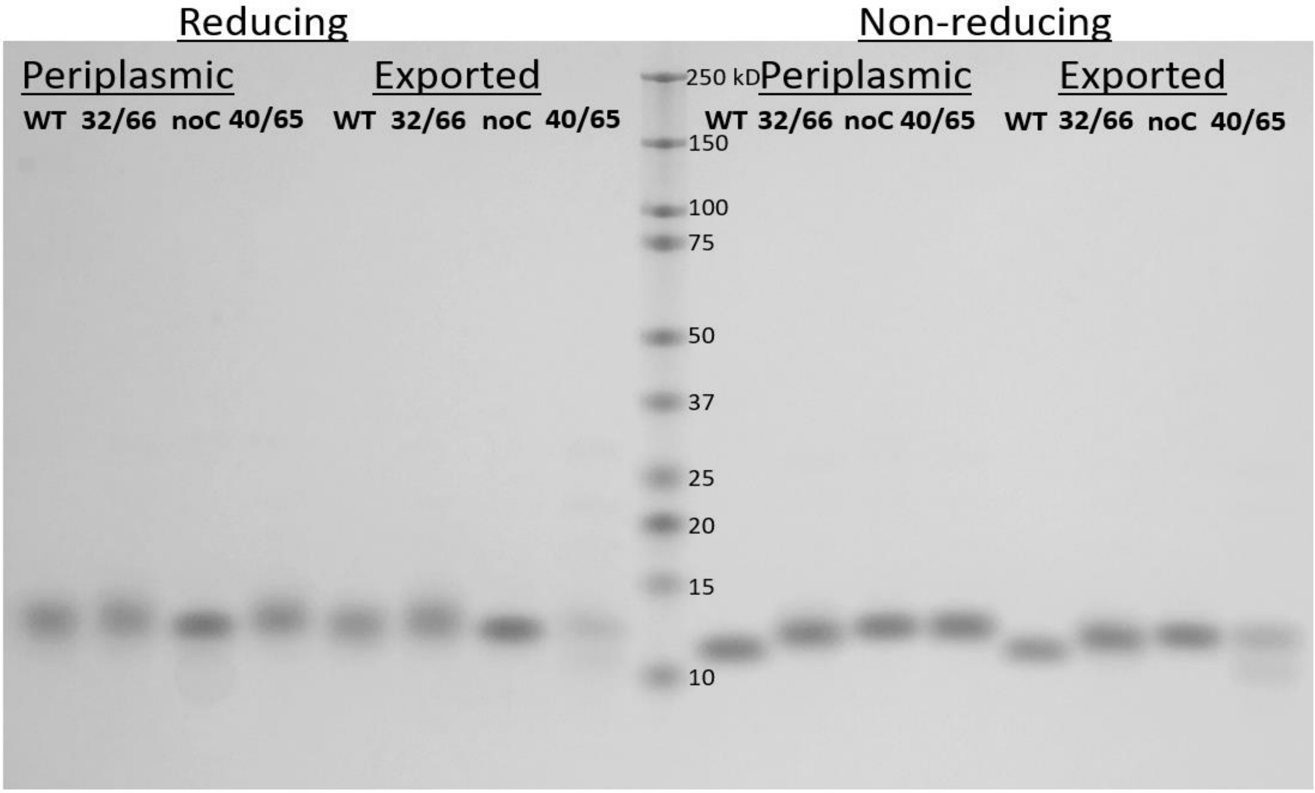


**Figure SI 6: Effect of disulfide bonds on HdeA variants' migration in SDS-PAGE.** Proteins were purified both from periplasmic extracts ("Periplasmic") and from the culture supernatants ("Exported"). Samples on the left side of the gel were treated with reducing agent (5 mM TCEP) during the denaturation step; samples on the right side were denatured without reduction.

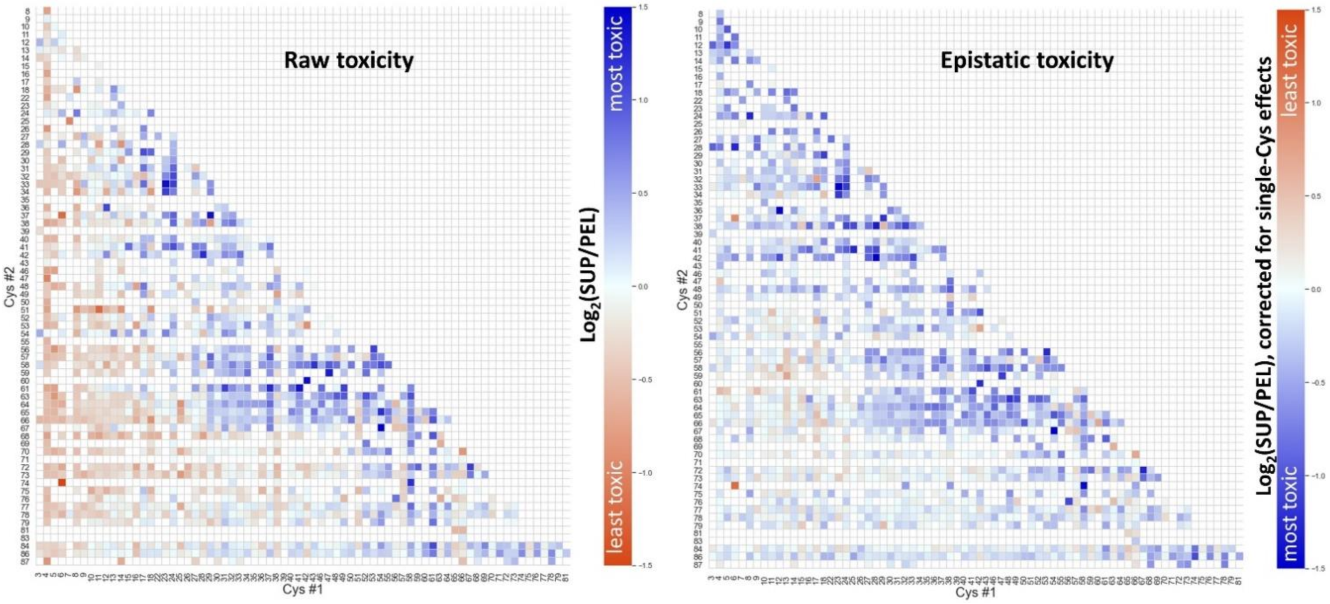


**Figure SI 7: Landscapes of allele toxicity of double-Cys HdeA variants.** *Left*: The raw average values of allele toxicity, as expressed by $\log_2$ ratios of the given allele's abundance in the culture supernatant ("SUP") vs. the cell pellet ("PEL"), showed that many Cys mutations in the disordered N-terminal region (approx. residues 1-14) resulted in rescue of toxicity compared to the noC variant (defined as 0 in this figure), but a clear cluster of highly toxic variants (approx. residues 29-49 x 58-66) was still evident. *Right*: When these raw values were corrected by dividing them by the product of toxicity effects of single-Cys variants at the corresponding positions, the signal from the N-terminal region largely disappeared, revealing a toxicity map quite similar to **Figure 3b**.

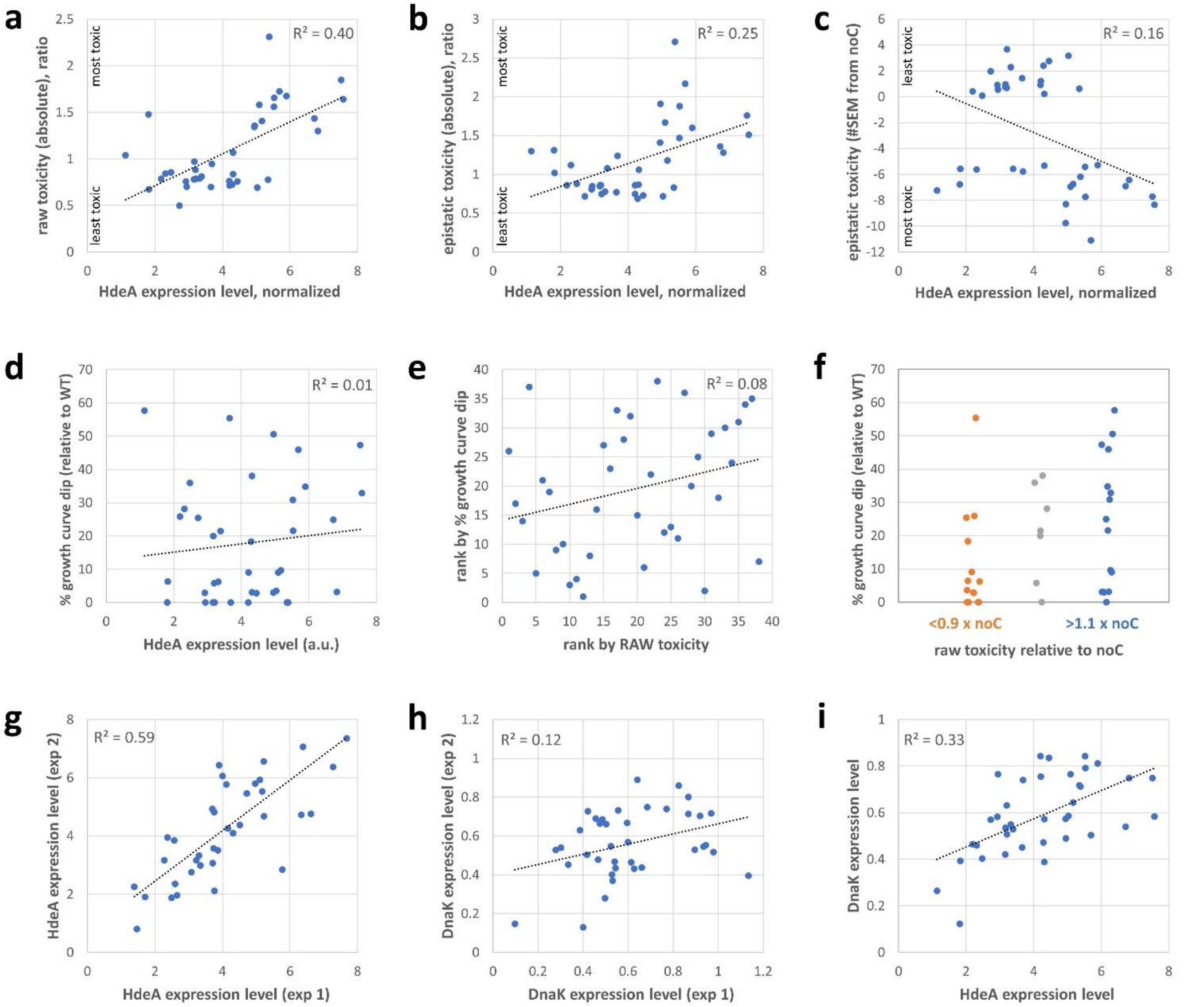


**Figure SI 8: Low-throughput characterization of expression level effects in 38 randomly chosen double-Cys HdeA variants.** Two independent expression experiments were conducted, with growth curves measured by $OD_{600}$ in 96-well format in sealed plates for 14 h and expression levels assayed by non-reducing SDS-PAGE from endpoint total culture samples from the same plates. HdeA and DnaK expression levels were internally normalized to the most intense band (37 kD) within each lane. Correlation between HdeA variant expression level and its raw allele toxicity **(a)** was much stronger than with epistatic toxicity **(b)**, especially if expressed as #SEM from noC **(c)**, the metric we use in the main text. Notably, there was no correlation between the magnitude of the dip in the growth curve and the HdeA expression level **(d)** and only a very weak correlation with the raw toxicity rank **(e)**. However, when variants were grouped by raw toxicity, the difference between group 1 (<90% of noC's toxicity) and group 3 (>110% of noC's toxicity) appeared significant by the one-tailed Mann-Whitney U test ($p = 0.025$). HdeA variant expression levels were reproducible between the two experiments **(f)**, but less so the DnaK expression levels **(g)**, which we attribute to the much fainter DnaK bands and therefore much higher noise in quantitating them. For all other panels, the expression levels from the two independent experiments were averaged to reduce noise. The correlation between HdeA and DnaK expression levels **(h)** should be considered as probably strong, given the high uncertainty in [DnaK] estimates **(g)**.

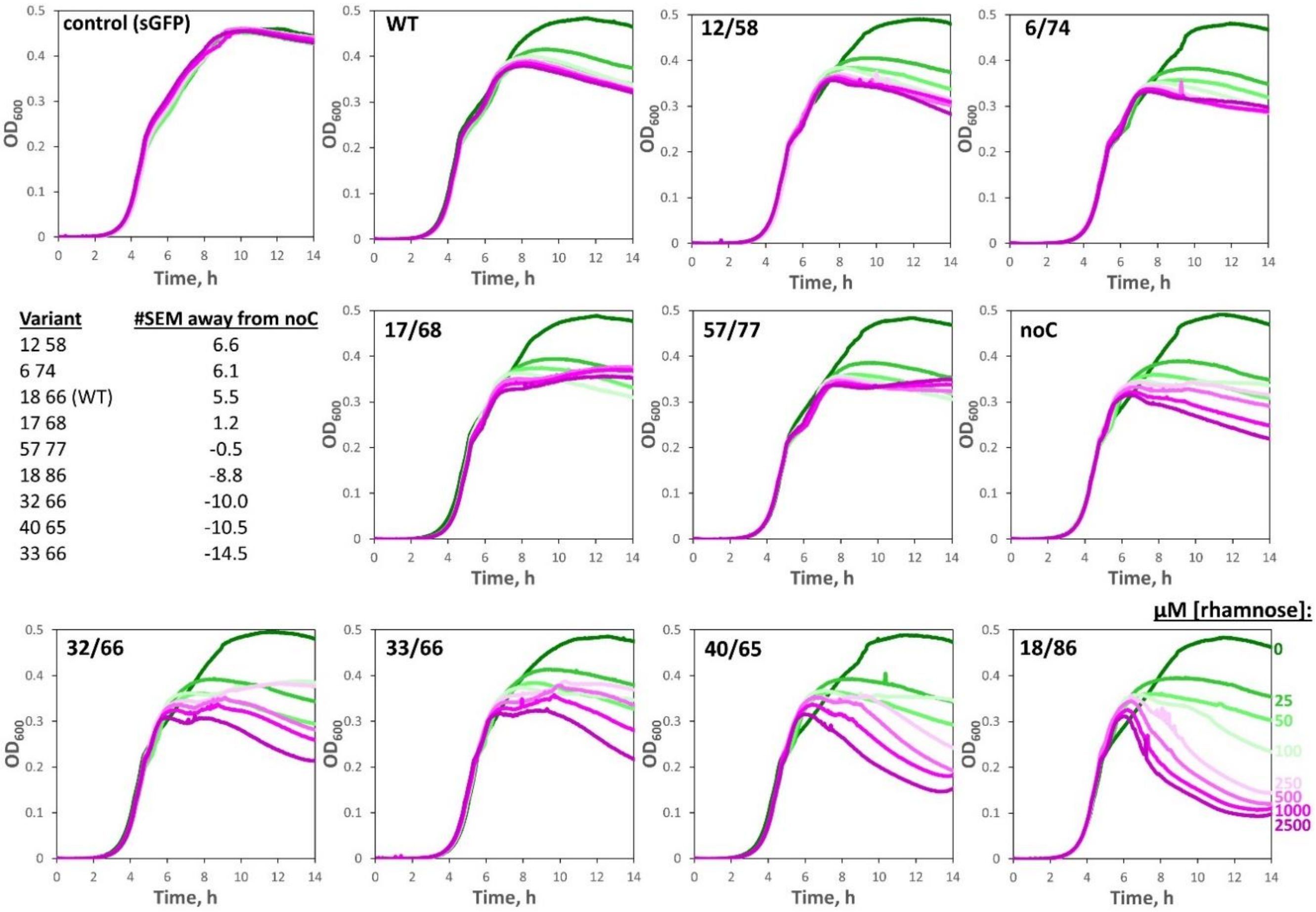


| Variant | #SEM away from noC |
|---|---|
| 12 58 | 6.6 |
| 6 74 | 6.1 |
| 18 66 (WT) | 5.5 |
| 17 68 | 1.2 |
| 57 77 | -0.5 |
| 18 86 | -8.8 |
| 32 66 | -10.0 |
| 40 65 | -10.5 |
| 33 66 | -14.5 |

**Figure SI 9: Dose-dependent growth curve shapes of select HdeA variants.** Growth curves of polyclonal transformant cultures in LB broth with antibiotics were monitored as in **Figure SI 2** with the indicated concentrations of inducer (rhamnose) present in the culture medium. While all HdeA variants showed some reduction of stationary-phase optical density when the proteins were expressed relative to no expression, the 12/58 and 6/74 variants were the only ones with WT-like traces, validating the results of the high-throughput assay in **Figure 3c**. The next-least-toxic variants, 17/68 and 57/77, showed a cusp upward in the stationary-phase growth curve at high induction. We did not investigate the origin of this effect, but it could be attributable, e.g., to cell lengthening induced by a mild heat shock – which would be consistent with pronounced cusps upward in the more toxic 32/66 and 40/65 variants at lower induction levels.

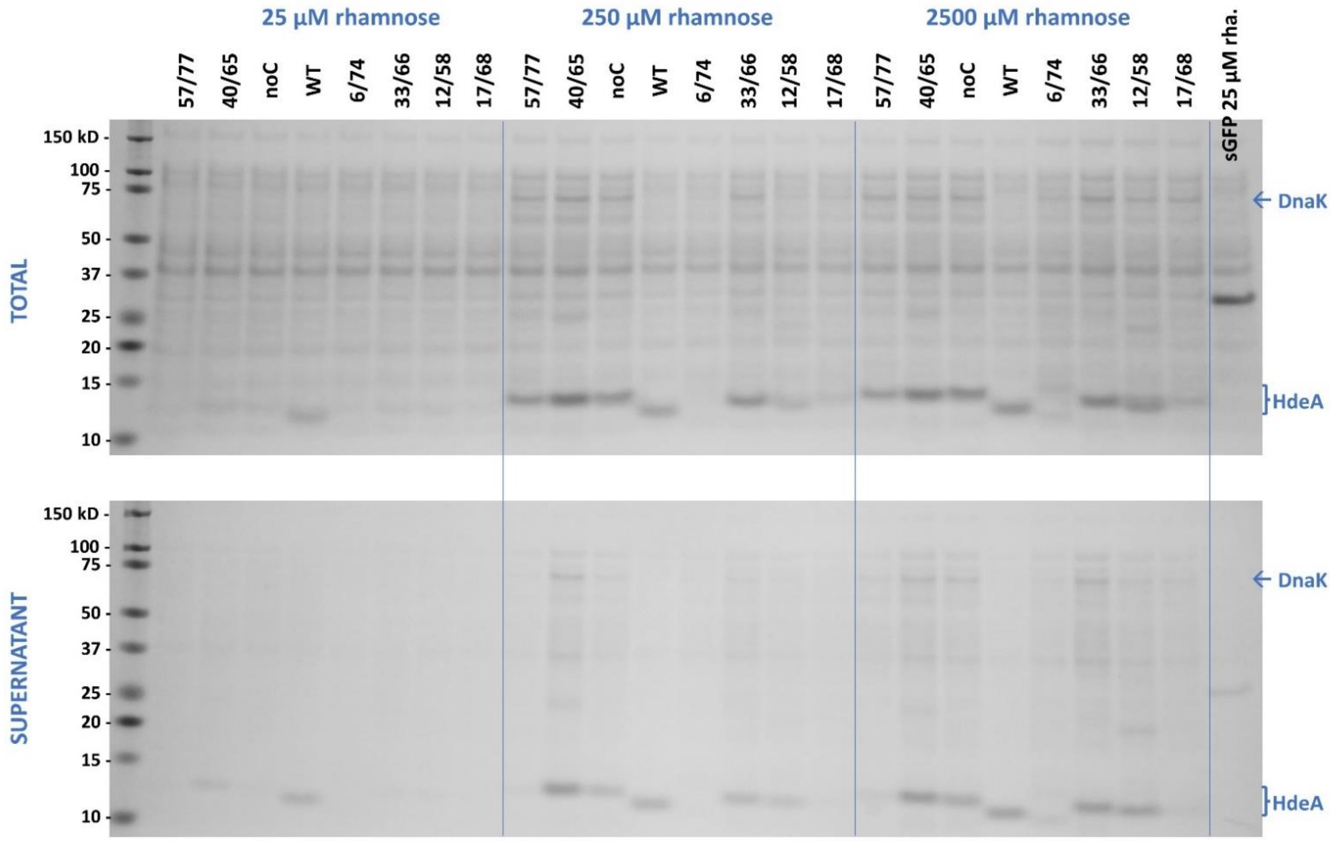


**Figure SI 10: Some non-toxic non-native double-Cys variants have exceptionally low levels of expression.** Non-reducing SDS-PAGE of total (*top gel*) and supernatant (*bottom gel*) samples from the endpoints of the growth curves in Figure SI 9, at 25, 250, and 2500 µM rhamnose, as indicated. The total culture samples showed greatly reduced expression of 6/74 and 17/68 relative to the WT, noC, or the toxic variants 40/65 and 33/66. Variant 12/58 also had a much reduced expression level at low or medium [inducer], but did accumulate to a significant level at the highest [inducer]. At the highest [inducer], however, both DnaK overexpression and cell lysis became evident even for that variant. (Note that culture spindown was not completely effective in this case, as evidenced by a trace of sGFP in the control lane on the bottom gel, as well as minor traces of cellular proteins in the supernatant of WT cultures; however, the much greater lysogenicity of the toxic variants is still visually apparent.) At the highest level of induction, even DnaK overexpression was visible in every HdeA variant except the WT. One unexpected observation was the near-total lack of export of the 57/77 variant from the overexpressing cells, despite significant accumulation comparable to the WT and the toxic mutants. We hypothesize that this variant may be retained in the cytoplasm of the cells, perhaps even in inclusion bodies, though other possibilities exist.

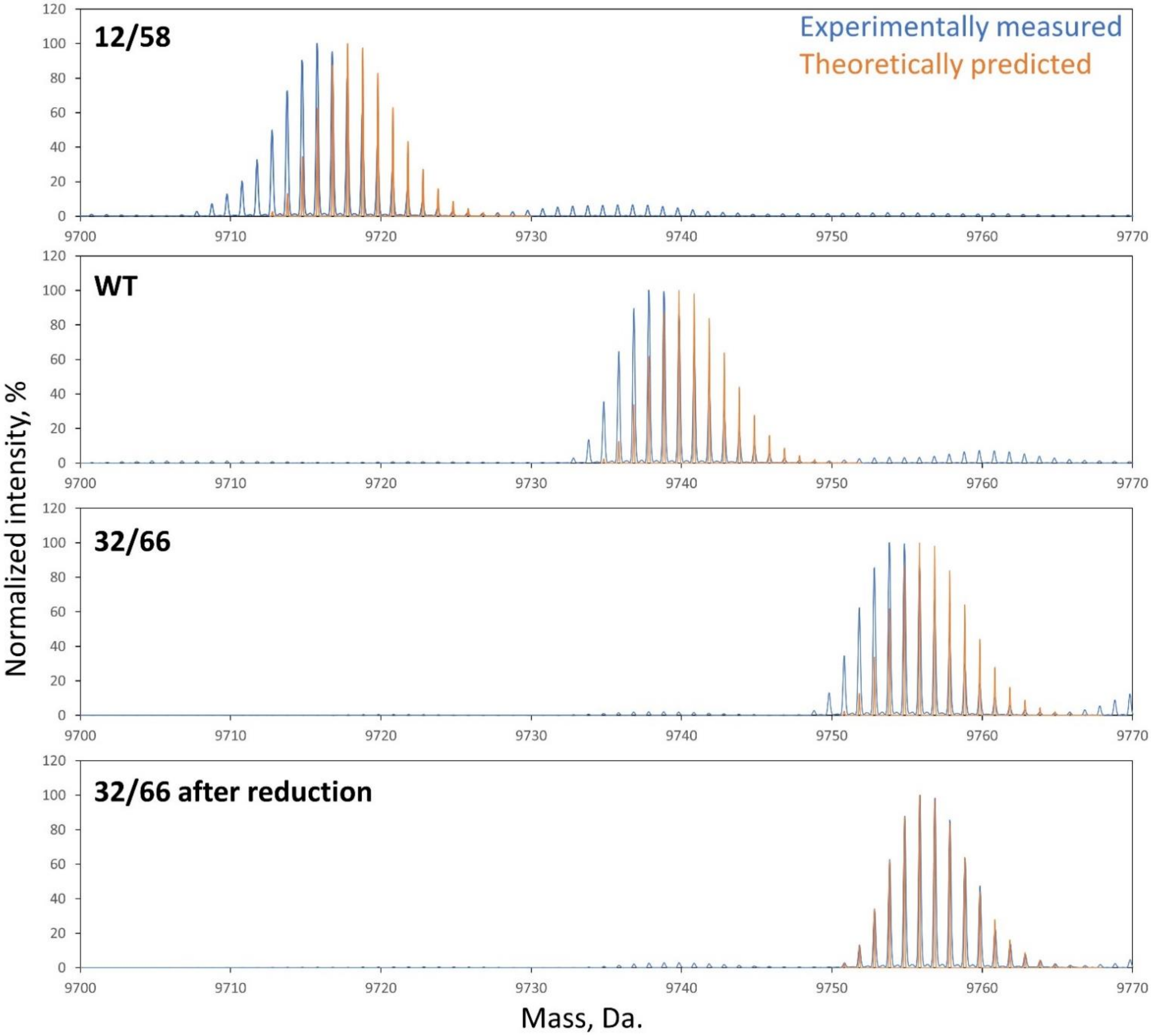


**Figure SI 11: Isotopically resolved electrospray mass spectrometry of intact HdeA variants confirms disulfide bond formation.** WT HdeA, as well as the 32/66 and 12/58 variants, were purified entirely in the disulfide-bonded state, as evidenced by the clear -2 Da. shift in the isotopic distribution compared to predicted values for the fully reduced proteins. The loss of two protons indicates oxidation of two -SH groups to form a disulfide. As a control (*bottom graph*), we carried out the same experiment with the same 32/66 sample after treating it with a reducing agent, which eliminated the -2 Da. shift. Minor peaks at +22 Da. were observed in all samples, attributable to a rare Na+ adduct in place of a proton.

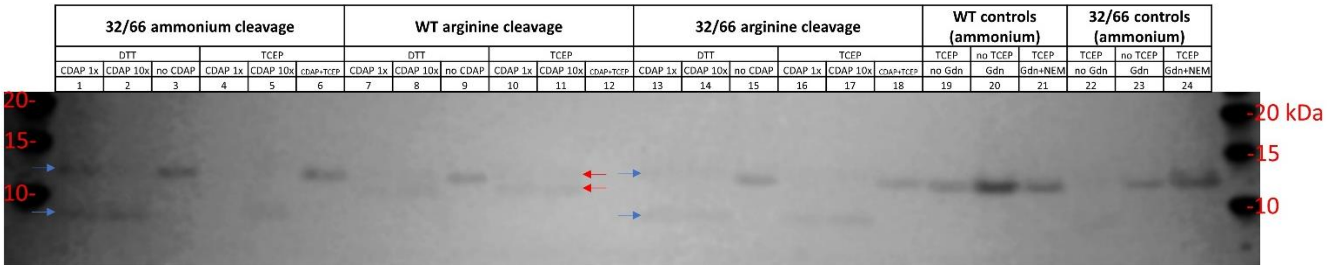


**Figure SI 12: Targeted cleavage of cyano-Cys containing HdeA by aminolysis.** Two variants (WT and 32/66) were used, since they yield fragments of distinct length upon Cys-specific cleavage. Prior to cyanylation, samples were reduced by either dithiothreitol (DTT) or tricarboxyethyl phosphine (TCEP), as indicated, with subsequent removal of the reducing agent by ultrafiltration. Either ammonium (3 M final concentration) or arginine (3 M final concentration) was used as the nucleophile for cleavage, as labeled. All ammonium-cleaved samples also contained 1 M (final concentration) of guanidinium chloride to further aid denaturation. All cleavage was at pH 9. Since samples containing high levels of guanidinium or arginine are not compatible with SDS-PAGE, all samples were desalted using C18 tips, with a wash step at 5% acetonitrile and elution in 50% acetonitrile, with 0.1% formic acid in all cases. The variable efficiency of elution and relatively low binding capacity of the C18 tips resulted in low and variable band intensities on the resulting gel; therefore, band intensities cannot be compared across lanes but should only be compared within each lane. Blue and red *arrows* indicate the intact (upper) and longest fragment (lower) bands for the 32/66 and WT (18/66) proteins, respectively. Several important results are evident by comparison of certain sets of lanes:

*Lanes 1 and 2* show that higher [CDAP] at the cyanylation step led to higher yield at the aminolysis step, suggesting that the limiting factor for aminolysis was the efficiency of cyanylation. “1x” = 1 mM.

*Lanes 3, 9, and 15* confirm that the pH 9 incubation with the nucleophiles did not by itself cause any detectable cleavage, meaning that cleavage was specific to cyano-Cys as expected.

*Lanes 2, 5, 11, 14, and 17* indicate that, at least at the higher [CDAP], overall reaction yield was very high. (Note that, since the cleavage fragment is shorter, it is expected to bind fewer Coomassie molecules per mole than the full-length protein, yet the intensities of the fragment bands in those lanes were higher than of the respective full-length bands.)

*Lanes 6, 12, and 18* indicate that presence of the reducing agent TCEP (added back after the ultrafiltration step) inhibited the cyanylation reaction, even though TCEP does not contain thiols (unlike DTT). Thus, removal of the reducing agent was important for reaction efficiency and is therefore preferable to the “one-pot” approach when high-efficiency cleavage is a priority.

*Lanes 19 and 22* show that the WT was not efficiently cleaved in the absence of initial denaturation (“no Gdn”), attributable to inaccessibility of the native disulfide bond in the folded protein to the reductant; by contrast, the 32/66 variant, with its predominantly molten structure, was still cleaved.

*Lanes 20 and 23* confirm that in the absence of initial reduction, no subsequent cleavage occurs; thus, only free thiols were modified by the cyanylating agent.

*Lanes 21 and 24* show that when thiol-blocking reagent (N-ethyl maleimide, NEM) was present during the initial reduction step with TCEP, no subsequent cleavage occurred, either; thus, Cys residues that are not oxidized but covalently modified by thiol-blocker were likewise not susceptible to cyanylation and cleavage.

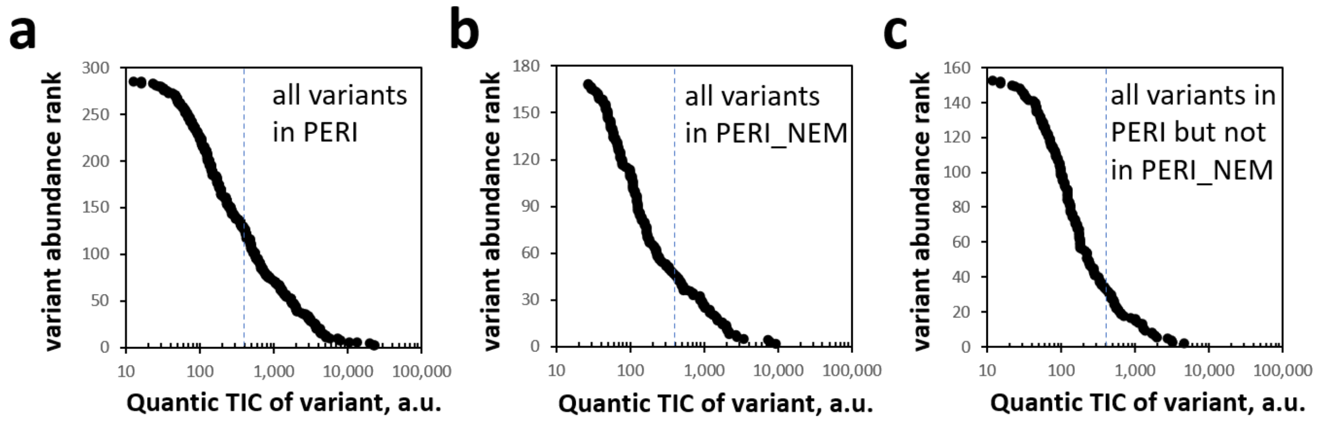


**Figure SI 13: Total ion current distributions for variants detected in Figure 4.** Peptide abundances are estimated by total ion current (TIC) using the Quantic tool of TPP6 Comet pipeline, based on intensities of top-6 MS/MS fragment ions. **(a)** Log-linear plot of all variants in PERI ranked by abundance revealed a sigmoidal distribution of abundance. **(b)** Same as **a** but for PERI_NEM. **(c)** Same as **a** but for variants found in PERI and not in PERI_NEM, showing that most were low-abundance variants (weak total ion currents). A blue dashed line at 400 a.u. is shown in all three plots for reference.

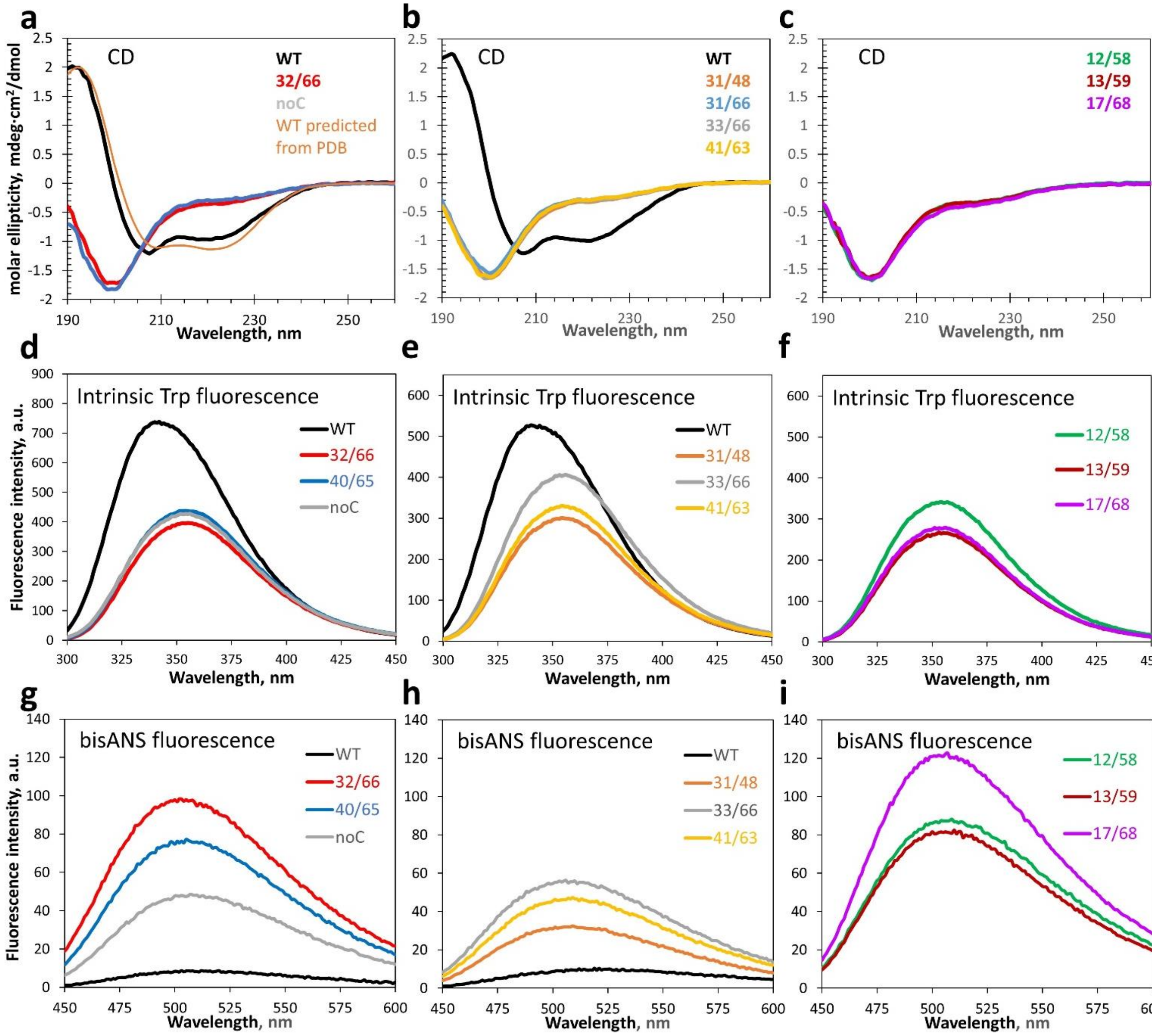


**Figure SI 14: Double-Cys variants do not rescue the native folded structure.** Panels **a**, **d**, and **g** are identical to **Figure 6c,a,d** and show the circular dichroism, intrinsic fluorescence, and bisANS fluorescence spectra for the WT, noC, 32/66, and 40/65 variants. Panels **b**, **e**, and **h** show the same data for the 31/48, 33/66, and 41/63 variants, as well as a separate sample of WT as an internal control. Panel **b** also includes the CD spectrum of the 31/66 variant. Panels **c**, **f**, and **i** show the same data for the 12/58, 13/59, and 17/68 variants.

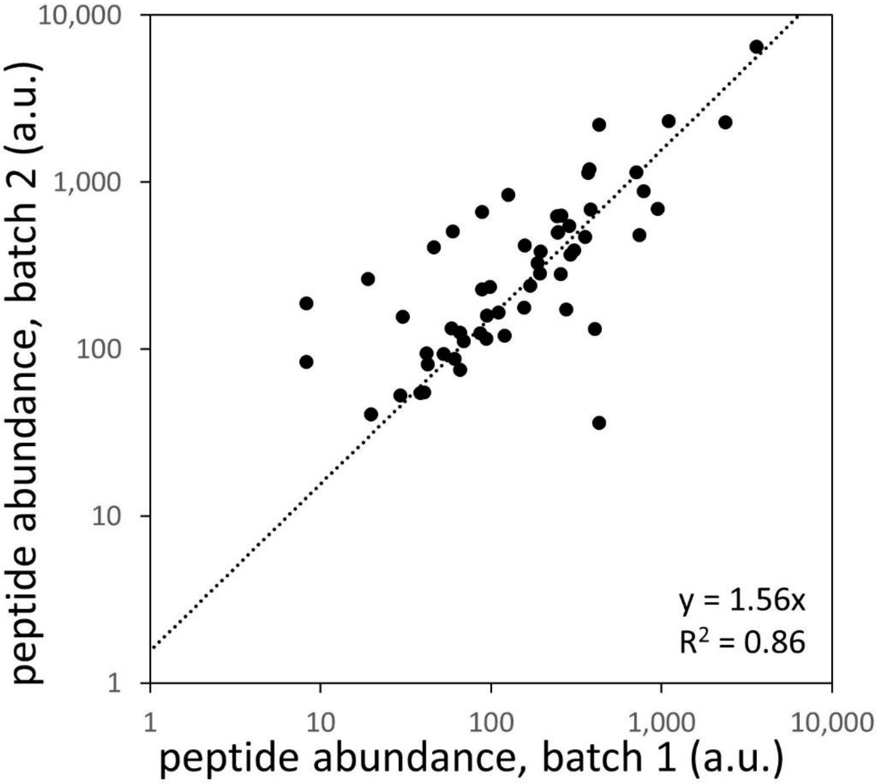


**Figure SI 15: Label-free peptide quantification by LC/MS/MS is reproducible.** Peptide abundances quantified by the Quantic tool of TPP6 Comet are plotted for the 56 variants identified in two separate batches (full biological replicates) of the multi-Cys scanning library. The true ratio of total protein concentrations injected onto LC/MS/MS for the two batches was 1.40, while the line of best fit returned a ratio of 1.56.

**Table SI 1: primer sequences**

fixed reverse primer: 5’-GAACTCAGAAGTGAAACG-3’

short mutagenic forward primers (mutated codon in **bold font**), 5'-3'

(a silent mutation in template is in red)

| | |
|---|---|
| noCssHdeAC3 | GCAGCCGAT**TGC**CAAAAAGCAGCT |
| noCssHdeAC4 | GCCGATGCG**TGC**AAAGCAGCTGAT |
| noCssHdeAC5 | GATGCGCAA**TGC**GCAGCTGATAAC |
| noCssHdeAC6 | GATGCGCAAAAA**TGC**GCTGATAAC |
| noCssHdeAC7 | CAAAAAGCA**TGC**GATAACAAAAAACCG |
| noCssHdeAC9 | GCAGCTGAT**TGC**AAAAAACCGGTC |
| noCssHdeAC10 | GCTGATAAC**TGC**AAACCGGTCAAC |
| noCssHdeAC11 | GATAACAAA**TGC**CCGGTCAACTCC |
| noCssHdeAC13 | CAAAAAACCG**TGC**AACTCCTGG |
| noCssHdeAC14 | CAAAAAACCGGTC**TGC**TCCTGGACC |
| noCssHdeAC15 | CCGGTCAAC**TGC**TGGACCAGT |
| noCssHdeAC17 | GTCAACTCCTGG**TGC**AGTGAAGATTTC |
| noCssHdeAC18 | TCCTGGACC**TGC**GAAGATTTCCTG |
| noCssHdeAC23 | GATTTCCTG**TGC**GTGGACGAATCC |
| noCssHdeAC24 | GATTTCCTGGCT**TGC**GACGAATCC |
| noCssHdeAC27 | GTGGACGAA**TGC**TTCCAGCCAACT |
| noCssHdeAC29 | GAATCCTTC**TGC**CCAACTGCA |
| noCssHdeAC31 | CTTCCAGCCA**TGC**GCAGTTGGT |
| noCssHdeAC32 | CAGCCAACT**TGC**GTTGGTTTTGCT |
| noCssHdeAC33 | CCAACTGCA**TGC**GGTTTTGCTGAAGCG |
| noCssHdeAC36 | GCAGTTGGTTTT**TGC**GAAGCGCTG |
| noCssHdeAC38 | GGTTTTGCTGAA**TGC**CTGAACAAC |
| noCssHdeAC40 | GAAGCGCTG**TGC**AACAAAGATAAACCA |
| noCssHdeAC41 | GAAGCGCTGAAC**TGC**AAAGATAAACCA |
| noCssHdeAC42 | GCGCTGAACAAC**TGC**GATAAACCA |
| noCssHdeAC48 | CCAGAAGAT**TGC**GTTTTAGATGTTCAG |
| noCssHdeAC49 | CCAGAAGATGCG**TGC**TTAGATGTTCAG |
| noCssHdeAC52 | GTTTTAGAT**TGC**CAGGGTATTGCAACC |
| noCssHdeAC53 | GTTTTAGATGTT**TGC**GGTATTGCAACC |
| noCssHdeAC56 | CAGGGTATT**TGC**ACCGTAACC |
| noCssHdeAC57 | GGTATTGCA**TGC**GTAACCCCA |
| noCssHdeAC58 | ATTGCAACC**TGC**ACCCCAGCTATC |
| noCssHdeAC59 | ATTGCAACCGTA**TGC**CCAGCTATC |
| noCssHdeAC61 | GTAACCCCA**TGC**ATCGTTCAGGCT |
| noCssHdeAC63 | CCAGCTATC**TGC**CAGGCTAGTACT |
| noCssHdeAC64 | GCTATCGTT**TGC**GCTAGTACTCAG |
| noCssHdeAC65 | GCTATCGTTCAG**TGC**AGTACTCAG |

| | |
|---|---|
| noCssHdeAC66 | GTTCAGGCT**TGC**ACTCAGGATAAACAAGCC |
| noCssHdeAC67 | CAGGCTAGT**TGC**CAGGATAAACAAGCC |
| noCssHdeAC68 | CAGGCTAGTACT**TGC**GATAAACAAGCC |
| noCssHdeAC70 | GCTAGTACTCAGGAT**TGC**CAAGCCAAC |
| noCssHdeAC71 | CAGGATAAA**TGC**GCCAACTTTAAAGAT |
| noCssHdeAC72 | CAGGATAAACAA**TGC**AACTTTAAAGAT |
| noCssHdeAC73 | GATAAACAAGCC**TGC**TTTAAAGATAAAGTT |
| noCssHdeAC75 | GCCAACTTT**TGC**GATAAAGTTAAAGGC |
| noCssHdeAC77 | GCCAACTTTAAAGAT**TGC**GTTAAAGGC |
| noCssHdeAC78 | CTTTAAAGATAAA**TGC**AAAGGCGAATGG |
| noCssHdeAC79 | CTTTAAAGATAAAGTT**TGC**GGCGAATGG |
| noCssHdeAC84 | GGCGAATGGGAC**TGC**ATTAAGAAAGAT |
| noCssHdeAC86 | GAATGGGACAAAATT**TGC**AAAGATATG |
| noCssHdeAC87 | GACAAAATTAAG**TGC**GATATGTAACACCATCAC |

**Table SI 2: HIC and toxicity values for 39 most abundant variants**

| variant | HIC peak, raw | HIC peak, corrected | #SEM from noC |
|---|---|---|---|
| 18/66 (WT) | 2.1 | 2.1 | 5.5 |
| 70/84 | 2.3 | 2.3 | -1.0 |
| 37/61 | 4.7 | 4.7 | -2.2 |
| 24/57 | 5.7 | 5.7 | -2.3 |
| 31/58 | 4.9 | 4.9 | -1.6 |
| 24/49 | 6.0 | 6.0 | -1.5 |
| 34/76 | 5.5 | 5.5 | -0.5 |
| 31/57 | 5.5 | 5.5 | -5.7 |
| 33/66 | 6.8 | 6.8 | -14.5 |
| 29/57 | 6.3 | 6.3 | -7.7 |
| 24/38 | 6.0 | 6.0 | -8.4 |
| 18/32 | 6.2 | 6.2 | -1.6 |
| 29/66 | 7.0 | 7.5 | -6.9 |
| 32/66 | 6.0 | 6.5 | -10.0 |
| 34/66 | 5.4 | 7.5 | -9.8 |
| 42/66 | 2.2 | 2.2 | -6.0 |
| 47/66 | 4.7 | 4.7 | -5.7 |
| 17/29 | 6.9 | 6.9 | -3.2 |
| 17/30 | 6.9 | 6.9 | -0.7 |
| 17/31 | 6.5 | 6.5 | -2.2 |
| 17/32 | 6.1 | 6.1 | 2.1 |
| 17/33 | 6.1 | 6.1 | -5.5 |

Color coding indicates which panel of Figure 7 of the main text shows the HIC trace.

**SI File 1: Comet (TPP6) parameters for middle peptides search**

```
# comet_version 2021.01 rev. 0
# Comet MS/MS search engine parameters file.
# Everything following the '#' symbol is treated as a comment.

database_name = /some/path/db.fasta
decoy_search = 0                    # 0=no (default), 1=concatenated search, 2=separate search
peff_format = 0                     # 0=no (normal fasta, default), 1=PEFF PSI-MOD, 2=PEFF Unimod
peff_obo = C:/TPP/conf/PSI-MOD.obo                        # path to PSI Mod or Unimod OBO file

num_threads = 0                     # 0=poll CPU to set num threads; else specify num threads directly (max 128)

#
# masses
#
peptide_mass_tolerance = 1.1
peptide_mass_units = 0              # 0=amu, 1=mmu, 2=ppm
mass_type_parent = 1                # 0=average masses, 1=monoisotopic masses
mass_type_fragment = 1              # 0=average masses, 1=monoisotopic masses
precursor_tolerance_type = 1        # 0=MH+ (default), 1=precursor m/z; only valid for amu/mmu tolerances
isotope_error = 3                   # 0=off, 1=0/1 (C13 error), 2=0/1/2, 3=0/1/2/3, 4=-8/-4/0/4/8 (for +4/+8 labeling)

#
# search enzyme
#
search_enzyme_number = 11           # choose from list at end of this params file
search_enzyme2_number = 0           # second enzyme; set to 0 if no second enzyme
num_enzyme_termini = 2              # 1 (semi-digested), 2 (fully digested, default), 8 C-term unspecific , 9 N-term
unspecific
allowed_missed_cleavage = 0         # maximum value is 5; for enzyme search

#
# Up to 9 variable modifications are supported
# format:  <mass> <residues> <0=variable/else binary> <max_mods_per_peptide> <term_distance> <n/c-term>
<required> <neutral_loss>
#     e.g. 79.966331 STY 0 3 -1 0 0 97.976896
#
variable_mod01 = 15.9949 M 0 3 -1 0 0 0.0
variable_mod02 = 26.02 C 1 1 0 2 1 0.0
variable_mod03 = 0.984 DE 0 1 -1 0 0 0.0
max_variable_mods_in_peptide = 3
require_variable_mod = 0

#
# fragment ions
#
# ion trap ms/ms:  1.0005 tolerance, 0.4 offset (mono masses), theoretical_fragment_ions = 1
```

```
# high res ms/ms:    0.02 tolerance, 0.0 offset (mono masses), theoretical_fragment_ions = 0,
spectrum_batch_size = 15000
#
fragment_bin_tol = 1.0005            # binning to use on fragment ions
fragment_bin_offset = 0.4            # offset position to start the binning (0.0 to 1.0)
theoretical_fragment_ions = 1         # 0=use flanking peaks, 1=M peak only
use_A_ions = 0
use_B_ions = 1
use_C_ions = 0
use_X_ions = 0
use_Y_ions = 1
use_Z_ions = 0
use_Z1_ions = 0
use_NL_ions = 1                      # 0=no, 1=yes to consider NH3/H2O neutral loss peaks

#
# output
#
output_sqtfile = 0                   # 0=no, 1=yes  write sqt file
output_txtfile = 0                   # 0=no, 1=yes  write tab-delimited txt file
output_pepxmlfile = 1                # 0=no, 1=yes  write pepXML file
output_mzidentmlfile = 0             # 0=no, 1=yes  write mzIdentML file
output_percolatorfile = 0            # 0=no, 1=yes  write Percolator pin file
print_expect_score = 1               # 0=no, 1=yes to replace Sp with expect in out & sqt
num_output_lines = 5                 # num peptide results to show

sample_enzyme_number = 11            # Sample enzyme which is possibly different than the one applied to the
search.
                                     # Used to calculate NTT & NMC in pepXML output (default=1 for trypsin).

#
# mzXML parameters
#
scan_range = 0 0                     # start and end scan range to search; either entry can be set independently
precursor_charge = 0 0               # precursor charge range to analyze; does not override any existing charge; 0 as
1st entry ignores parameter
override_charge = 0                  # 0=no, 1=override precursor charge states, 2=ignore precursor charges outside
precursor_charge range, 3=see online
ms_level = 2                         # MS level to analyze, valid are levels 2 (default) or 3
activation_method = ALL              # activation method; used if activation method set; allowed ALL, CID, ECD,
ETD, ETD+SA, PQD, HCD, IRMPD, SID

#
# misc parameters
#
digest_mass_range = 500.0 10000.0      # MH+ peptide mass range to analyze
peptide_length_range = 5 63            # minimum and maximum peptide length to analyze (default 1 63; max
length 63)
```

num_results = 100 # number of search hits to store internally
max_duplicate_proteins = -1 # maximum number of additional duplicate protein names to report for each peptide ID; -1 reports all duplicates
max_fragment_charge = 5 # set maximum fragment charge state to analyze (allowed max 5)
max_precursor_charge = 9 # set maximum precursor charge state to analyze (allowed max 9)
nucleotide_reading_frame = 0 # 0=proteinDB, 1-6, 7=forward three, 8=reverse three, 9=all six
clip_nterm_methionine = 0 # 0=leave sequences as-is; 1=also consider sequence w/o N-term methionine
spectrum_batch_size = 15000 # max. # of spectra to search at a time; 0 to search the entire scan range in one loop
decoy_prefix = DECOY_ # decoy entries are denoted by this string which is pre-pended to each protein accession
equal_I_and_L = 1 # 0=treat I and L as different; 1=treat I and L as same
output_suffix = # add a suffix to output base names i.e. suffix "-C" generates base-C.pep.xml from base.mzXML input
mass_offsets = # one or more mass offsets to search (values substracted from deconvoluted precursor mass)
precursor_NL_ions = # one or more precursor neutral loss masses, will be added to xcorr analysis

#
### spectral processing
#
minimum_peaks = 10 # required minimum number of peaks in spectrum to search (default 10)
minimum_intensity = 0 # minimum intensity value to read in
remove_precursor_peak = 0 # 0=no, 1=yes, 2=all charge reduced precursor peaks (for ETD), 3=phosphate neutral loss peaks
remove_precursor_tolerance = 1.5 # +- Da tolerance for precursor removal
clear_mz_range = 0.0 0.0 # for iTRAQ/TMT type data; will clear out all peaks in the specified m/z range

#
### additional modifications
#

add_Cterm_peptide = 0.0
add_Nterm_peptide = 0.0
add_Cterm_protein = 0.0
add_Nterm_protein = 0.0

add_G_glycine = 0.0000 # added to G - avg. 57.0513, mono. 57.02146
add_A_alanine = 0.0000 # added to A - avg. 71.0779, mono. 71.03711
add_S_serine = 0.0000 # added to S - avg. 87.0773, mono. 87.03203
add_P_proline = 0.0000 # added to P - avg. 97.1152, mono. 97.05276
add_V_valine = 0.0000 # added to V - avg. 99.1311, mono. 99.06841
add_T_threonine = 0.0000 # added to T - avg. 101.1038, mono. 101.04768
add_C_cysteine = 0.0000 # added to C - avg. 103.1429, mono. 103.00918
add_L_leucine = 0.0000 # added to L - avg. 113.1576, mono. 113.08406
add_I_isoleucine = 0.0000 # added to I - avg. 113.1576, mono. 113.08406
add_N_asparagine = 0.0000 # added to N - avg. 114.1026, mono. 114.04293
add_D_aspartic_acid = 0.0000 # added to D - avg. 115.0874, mono. 115.02694

```
add_Q_glutamine = 0.0000             # added to Q - avg. 128.1292, mono. 128.05858
add_K_lysine = 0.0000                # added to K - avg. 128.1723, mono. 128.09496
add_E_glutamic_acid = 0.0000         # added to E - avg. 129.1140, mono. 129.04259
add_M_methionine = 0.0000            # added to M - avg. 131.1961, mono. 131.04048
add_H_histidine = 0.0000             # added to H - avg. 137.1393, mono. 137.05891
add_F_phenylalanine = 0.0000         # added to F - avg. 147.1739, mono. 147.06841
add_U_selenocysteine = 0.0000        # added to U - avg. 150.0379, mono. 150.95363
add_R_arginine = 0.0000              # added to R - avg. 156.1857, mono. 156.10111
add_Y_tyrosine = 0.0000              # added to Y - avg. 163.0633, mono. 163.06333
add_W_tryptophan = 0.0000            # added to W - avg. 186.0793, mono. 186.07931
add_O_pyrrolysine = 0.0000           # added to O - avg. 237.2982, mono  237.14773
add_B_user_amino_acid = 0.0000       # added to B - avg.   0.0000, mono.   0.00000
add_J_user_amino_acid = 0.0000       # added to J - avg.   0.0000, mono.   0.00000
add_X_user_amino_acid = 0.0000       # added to X - avg.   0.0000, mono.   0.00000
add_Z_user_amino_acid = 0.0000       # added to Z - avg.   0.0000, mono.   0.00000

#
# COMET_ENZYME_INFO _must_ be at the end of this parameters file
#
[COMET_ENZYME_INFO]
0.  Cut_everywhere         0      -           -
1.  Trypsin                1      KR          P
2.  Trypsin/P              1      KR          -
3.  Lys_C                  1      K           P
4.  Lys_N                  0      K           -
5.  Arg_C                  1      R           P
6.  Asp_N                  0      D           -
7.  CNBr                   1      M           -
8.  Glu_C                  1      DE          P
9.  PepsinA                1      FL          P
10. Chymotrypsin           1      FWYL        P
11. CDAP                   0      C           -
12. No_cut                 1      @           @
```

**SI File 2: Comet (TPP6) parameters for N-terminal peptides search**

```
# comet_version 2021.01 rev. 0
# Comet MS/MS search engine parameters file.
# Everything following the '#' symbol is treated as a comment.

database_name = /some/path/db.fasta
decoy_search = 0                    # 0=no (default), 1=concatenated search, 2=separate search
peff_format = 0                     # 0=no (normal fasta, default), 1=PEFF PSI-MOD, 2=PEFF Unimod
peff_obo = C:/TPP/conf/PSI-MOD.obo                    # path to PSI Mod or Unimod OBO file

num_threads = 0                     # 0=poll CPU to set num threads; else specify num threads directly (max 128)

#
# masses
#
peptide_mass_tolerance = 1.1
peptide_mass_units = 0              # 0=amu, 1=mmu, 2=ppm
mass_type_parent = 1                # 0=average masses, 1=monoisotopic masses
mass_type_fragment = 1              # 0=average masses, 1=monoisotopic masses
precursor_tolerance_type = 1        # 0=MH+ (default), 1=precursor m/z; only valid for amu/mmu tolerances
isotope_error = 3                   # 0=off, 1=0/1 (C13 error), 2=0/1/2, 3=0/1/2/3, 4=-8/-4/0/4/8 (for +4/+8 labeling)

#
# search enzyme
#
search_enzyme_number = 12           # choose from list at end of this params file
search_enzyme2_number = 0           # second enzyme; set to 0 if no second enzyme
num_enzyme_termini = 2              # 1 (semi-digested), 2 (fully digested, default), 8 C-term unspecific , 9 N-term
unspecific
allowed_missed_cleavage = 0         # maximum value is 5; for enzyme search

#
# Up to 9 variable modifications are supported
# format:  <mass> <residues> <0=variable/else binary> <max_mods_per_peptide> <term_distance> <n/c-term>
<required> <neutral_loss>
#     e.g. 79.966331 STY 0 3 -1 0 0 97.976896
#
variable_mod01 = 15.9949 M 0 3 -1 0 0 0.0
variable_mod02 = 0.984 DE 0 1 -1 0 0 0.0
max_variable_mods_in_peptide = 3
require_variable_mod = 0

#
# fragment ions
#
# ion trap ms/ms:  1.0005 tolerance, 0.4 offset (mono masses), theoretical_fragment_ions = 1
```

# high res ms/ms:    0.02 tolerance, 0.0 offset (mono masses), theoretical_fragment_ions = 0, spectrum_batch_size = 15000
#
fragment_bin_tol = 1.0005             # binning to use on fragment ions
fragment_bin_offset = 0.4             # offset position to start the binning (0.0 to 1.0)
theoretical_fragment_ions = 1         # 0=use flanking peaks, 1=M peak only
use_A_ions = 0
use_B_ions = 1
use_C_ions = 0
use_X_ions = 0
use_Y_ions = 1
use_Z_ions = 0
use_Z1_ions = 0
use_NL_ions = 1                       # 0=no, 1=yes to consider NH3/H2O neutral loss peaks

#
### output
#
output_sqtfile = 0                    # 0=no, 1=yes  write sqt file
output_txtfile = 0                    # 0=no, 1=yes  write tab-delimited txt file
output_pepxmlfile = 1                 # 0=no, 1=yes  write pepXML file
output_mzidentmlfile = 0              # 0=no, 1=yes  write mzIdentML file
output_percolatorfile = 0             # 0=no, 1=yes  write Percolator pin file
print_expect_score = 1                # 0=no, 1=yes to replace Sp with expect in out & sqt
num_output_lines = 5                  # num peptide results to show

sample_enzyme_number = 11             # Sample enzyme which is possibly different than the one applied to the search.
                                      # Used to calculate NTT & NMC in pepXML output (default=1 for trypsin).

#
### mzXML parameters
#
scan_range = 0 0                      # start and end scan range to search; either entry can be set independently
precursor_charge = 0 0                # precursor charge range to analyze; does not override any existing charge; 0 as 1st entry ignores parameter
override_charge = 0                   # 0=no, 1=override precursor charge states, 2=ignore precursor charges outside precursor_charge range, 3=see online
ms_level = 2                          # MS level to analyze, valid are levels 2 (default) or 3
activation_method = ALL               # activation method; used if activation method set; allowed ALL, CID, ECD, ETD, ETD+SA, PQD, HCD, IRMPD, SID

#
### misc parameters
#
digest_mass_range = 500.0 10000.0     # MH+ peptide mass range to analyze
peptide_length_range = 5 63           # minimum and maximum peptide length to analyze (default 1 63; max length 63)

```
num_results = 100                    # number of search hits to store internally
max_duplicate_proteins = -1          # maximum number of additional duplicate protein names to report for each
peptide ID; -1 reports all duplicates
max_fragment_charge = 5              # set maximum fragment charge state to analyze (allowed max 5)
max_precursor_charge = 9             # set maximum precursor charge state to analyze (allowed max 9)
nucleotide_reading_frame = 0         # 0=proteinDB, 1-6, 7=forward three, 8=reverse three, 9=all six
clip_nterm_methionine = 0            # 0=leave sequences as-is; 1=also consider sequence w/o N-term methionine
spectrum_batch_size = 15000          # max. # of spectra to search at a time; 0 to search the entire scan range in
one loop
decoy_prefix = DECOY_                # decoy entries are denoted by this string which is pre-pended to each protein
accession
equal_I_and_L = 1                    # 0=treat I and L as different; 1=treat I and L as same
output_suffix =                      # add a suffix to output base names i.e. suffix "-C" generates base-C.pep.xml from
base.mzXML input
mass_offsets =                       # one or more mass offsets to search (values substracted from deconvoluted
precursor mass)
precursor_NL_ions =                  # one or more precursor neutral loss masses, will be added to xcorr analysis

#
# spectral processing
#
minimum_peaks = 10                   # required minimum number of peaks in spectrum to search (default 10)
minimum_intensity = 0                # minimum intensity value to read in
remove_precursor_peak = 0            # 0=no, 1=yes, 2=all charge reduced precursor peaks (for ETD), 3=phosphate
neutral loss peaks
remove_precursor_tolerance = 1.5     # +- Da tolerance for precursor removal
clear_mz_range = 0.0 0.0             # for iTRAQ/TMT type data; will clear out all peaks in the specified m/z range

#
# additional modifications
#

add_Cterm_peptide = 0.0
add_Nterm_peptide = 0.0
add_Cterm_protein = 0.0
add_Nterm_protein = 0.0

add_G_glycine = 0.0000               # added to G - avg.  57.0513, mono.  57.02146
add_A_alanine = 0.0000               # added to A - avg.  71.0779, mono.  71.03711
add_S_serine = 0.0000                # added to S - avg.  87.0773, mono.  87.03203
add_P_proline = 0.0000               # added to P - avg.  97.1152, mono.  97.05276
add_V_valine = 0.0000                # added to V - avg.  99.1311, mono.  99.06841
add_T_threonine = 0.0000             # added to T - avg. 101.1038, mono. 101.04768
add_C_cysteine = 0.0000              # added to C - avg. 103.1429, mono. 103.00918
add_L_leucine = 0.0000               # added to L - avg. 113.1576, mono. 113.08406
add_I_isoleucine = 0.0000            # added to I - avg. 113.1576, mono. 113.08406
add_N_asparagine = 0.0000            # added to N - avg. 114.1026, mono. 114.04293
add_D_aspartic_acid = 0.0000         # added to D - avg. 115.0874, mono. 115.02694
```

```
add_Q_glutamine = 0.0000             # added to Q - avg. 128.1292, mono. 128.05858
add_K_lysine = 0.0000                # added to K - avg. 128.1723, mono. 128.09496
add_E_glutamic_acid = 0.0000         # added to E - avg. 129.1140, mono. 129.04259
add_M_methionine = 0.0000            # added to M - avg. 131.1961, mono. 131.04048
add_H_histidine = 0.0000             # added to H - avg. 137.1393, mono. 137.05891
add_F_phenylalanine = 0.0000         # added to F - avg. 147.1739, mono. 147.06841
add_U_selenocysteine = 0.0000        # added to U - avg. 150.0379, mono. 150.95363
add_R_arginine = 0.0000              # added to R - avg. 156.1857, mono. 156.10111
add_Y_tyrosine = 0.0000              # added to Y - avg. 163.0633, mono. 163.06333
add_W_tryptophan = 0.0000            # added to W - avg. 186.0793, mono. 186.07931
add_O_pyrrolysine = 0.0000           # added to O - avg. 237.2982, mono  237.14773
add_B_user_amino_acid = 0.0000       # added to B - avg.   0.0000, mono.   0.00000
add_J_user_amino_acid = 0.0000       # added to J - avg.   0.0000, mono.   0.00000
add_X_user_amino_acid = 0.0000       # added to X - avg.   0.0000, mono.   0.00000
add_Z_user_amino_acid = 0.0000       # added to Z - avg.   0.0000, mono.   0.00000

#
# COMET_ENZYME_INFO _must_ be at the end of this parameters file
#
[COMET_ENZYME_INFO]
0.  Cut_everywhere         0      -           -
1.  Trypsin                1      KR          P
2.  Trypsin/P              1      KR          -
3.  Lys_C                  1      K           P
4.  Lys_N                  0      K           -
5.  Arg_C                  1      R           P
6.  Asp_N                  0      D           -
7.  CNBr                   1      M           -
8.  Glu_C                  1      DE          P
9.  PepsinA                1      FL          P
10. Chymotrypsin           1      FWYL        P
11. CDAP                   0      C           -
12. No_cut                 1      @           @
```
